# C11orf95-RELA Reprograms 3D Epigenome in Supratentorial Ependymoma

**DOI:** 10.1101/2020.06.04.135079

**Authors:** Jacqueline Jufen Zhu, Nathaniel Jillette, Xiao-Nan Li, Albert Wu Cheng, Ching C Lau

## Abstract

Supratentorial ependymoma (ST-EPN) is a type of malignant brain tumor mainly seen in children. Since 2014, it has been known that an intrachromosomal fusion *C11orf95-RELA* is an oncogenic driver in ST-EPN^1^ but the molecular mechanisms of oncogenesis are unclear. Here we show that the C11orf95 component of the fusion protein dictates DNA binding activity while the RELA component is required for driving the expression of ependymoma-associated genes. Epigenomic characterizations using ChIP-seq and HiChIP approaches reveal that C11orf95-RELA modulates chromatin states and mediates chromatin interactions, leading to transcriptional reprogramming in ependymoma cells. Our findings provide important characterization of the molecular underpinning of *C11orf95-RELA* fusion and shed light on potential therapeutic targets for *C11orf95-RELA* subtype ependymoma.

---

**Ependymoma is the third most common malignant brain tumor in children. It can be classified into three groups based on location of the tumor: supratentorial, infratentorial and spinal. Currently there is no effective chemotherapy identified and treatment is limited to surgery with or without adjuvant radiation therapy. Because a significant portion of ependymoma is found in young children, using radiation therapy in such patients is not desirable due to the detrimental effects on the developing brain. Thus, there is an urgent need to develop targeted therapy based on the underlying biology. *C11orf95-RELA* fusion was found to be the most recurrent structural variation in approximately 70% of supratentorial ependymomas (ST-EPN)**^**1**^**. Neural stem cells transformed by C11orf95-RELA were able to form brain tumor in mice**^**1**,**2**^**. However, how C11orf95-RELA functions in tumorigenesis at the molecular level is largely unknown. We present results here showing that contrary to the hypothesis that C11orf95-RELA ependymoma is driven by the RELA component of the fusion, binding affinity of the fusion protein to DNA is dictated by the C11orf95 component. The contribution of the RELA component is to stabilize the DNA binding of the fusion protein and provide its activation domain to drive expression of the target genes.**

## C11orf95-RELA is a novel transcription factor that recognizes a specific DNA motif dictated by the *C11orf95* fragment

*C11orf95-RELA*^*fus1*^ was identified as the most frequently occurring fusion subtype in supratentorial ependymoma with *C11orf95-RELA* fusion (ST-EPN-RELA), containing almost the entire *RELA* gene except the first two codons fused to a truncated *C11orf95* gene fragment (*C11orf95*^*fus1*^) harboring two and half exons out of five exons of the full-size gene^1^. Unlike RELA, which is a well-known transcription factor subunit of NF-κB, C11orf95 is an uncharacterized protein. In ST-EPN-RELA cells, RELA was found to be constitutively localized in the nucleus after fusing with C11orf95^1^, leading to the hypothesis that C11orf95-RELA is a novel transcription factor. Interestingly, our study showed that C11orf95^fus1^ exclusively localized to nucleus when overexpressed in HEK293T cells (Extended Data Fig. 1), implying its DNA binding potential. However, there was previously no direct evidence for the binding of C11orf95^fus1^ and C11orf95-RELA^fus1^ to DNA. To address this question and further decipher the regulatory mechanisms of the fusion protein, we engineered HEK293T cell lines to express C11orf95-RELA^fus1^ (G16-4), C11orf95^fus1^ fragment (G16-3), or the RELA fragment (G16-2) in order to study the contribution of the fusion protein and each of the partners to epigenetic program and transcriptional regulation. These transgenes are tagged with the HA-epitope and are under the control of a doxycycline-inducible TetO promoter (Extended Data Figs. 2a, b). The HA tag facilitates our molecular experimentations such as Western blot and ChIP, despite the lack of good quality antibodies against the fusion and C11orf95 proteins. We then conducted ChIP-seq using HA antibody to profile genome-wide chromatin bindings of RELA, C11orf95^fus1^ and C11orf95-RELA^fus1^ in G16-2, G16-3 and G16-4 cells, respectively. Surprisingly, 32,152 and 38,428 binding peaks were identified for C11orf95-RELA^fus1^ and C11orf95^fus1^, respectively, with 16,829 overlapping (Fig. 1a). In contrast, RELA ChIP-seq only identified 111 binding sites in total. Considering RELA is normally inactive without cellular stimulation, we treated G16-2 cells with tumor necrosis factor (TNF) for six hours after doxycycline induction of RELA expression, in order to capture RELA bindings in its active state. As a result, 1542 binding peaks were identified, which was still far less than the fusion or C11orf95 bindings. Of these RELA peaks after TNF stimulation, less than 25% overlapped with C11orf95-RELA^fus1^ peaks (Fig. 1a). These results indicate that C11orf95^fus1^ contributes to the majority of DNA binding of the fusion protein.

**Fig. 1.**
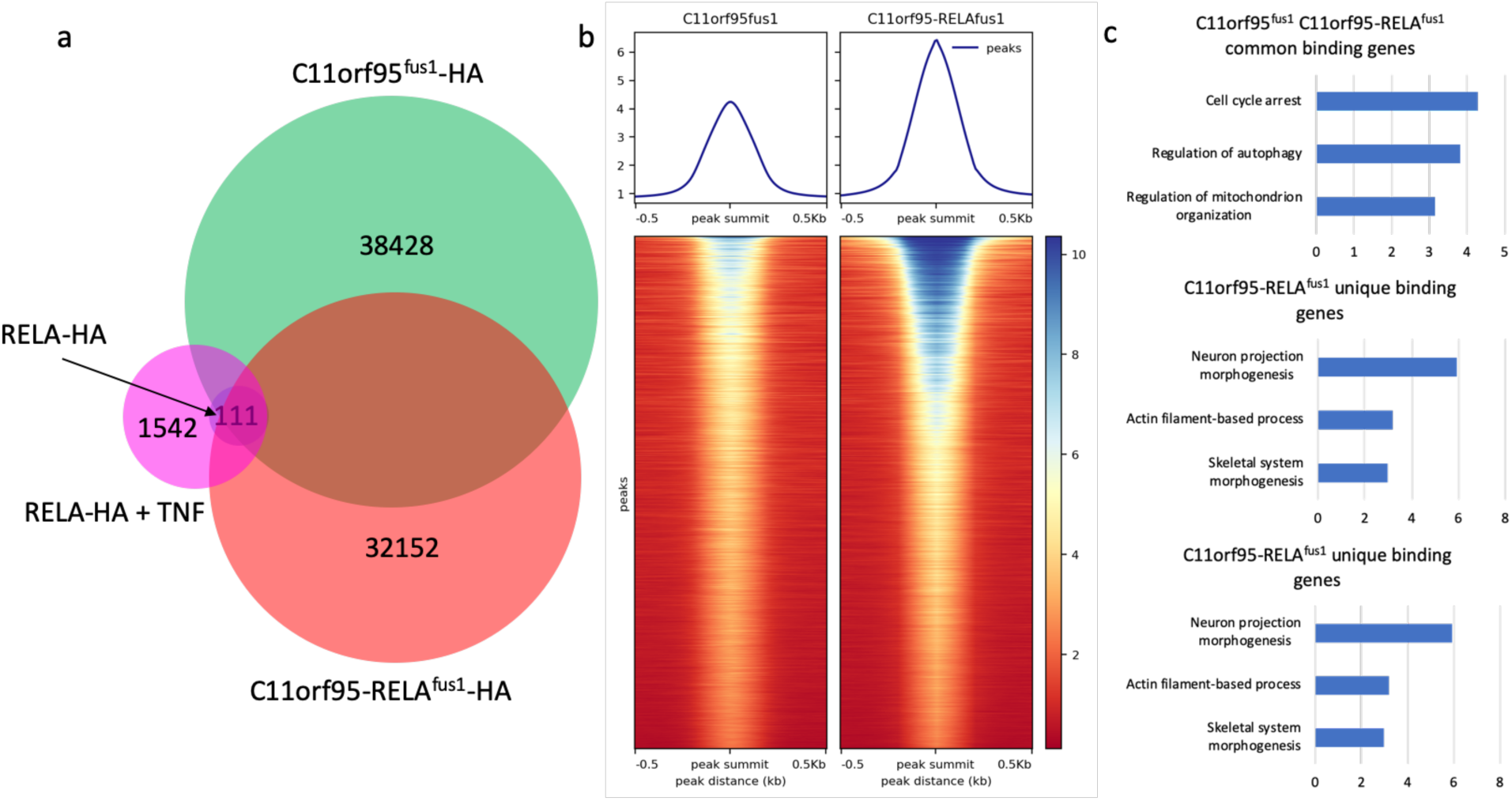
a. Venn diagram of ChIP-seq peaks. The DNA pull-down were done by RELA-HA (blue, the smallest circle) in G16-2 cells, RELA-HA after TNF treatment (magenta) in G16-2 cells, C11orf95^fus1^-HA (green) in G16-3 cells and C11orf95-RELAf^us1^-HA (red) in G16-4 cells. The numbers indicate the total peaks identified in each cohort. The Venn diagram is not dawn to scale. b. Read densities over ChIP-seq peaks of C11orf95^fus1^ and C11orf95-RELA^fus1^. Using peak summit as the center, the normalized mapped reads over a 1kb region around the peak were plotted for a total of 16,829 shared peaks of C11orf95^fus1^ and C11orf95-RELA^fus1^. c. Gene ontology analysis on C11orf95^fus1^ and C11orf95-RELA^fus1^ common binding genes, C11orf95^fus1^ unique binding genes and C11orf95-RELA^fus1^ unique binding genes. The top three ranked biological processes according to q-value are shown.

Although both C11orf95^fus1^ and C11orf95-RELA^fus1^ can bind to DNA, their binding targets were not completely identical. Other than the common peaks, there were also specific peaks for each of the two proteins (Fig. 1a). Among the 16,829 shared peaks, C11orf95-RELA^fus1^ had significantly higher signal densities than C11orf95^fus1^ after sequencing read normalization (Fig. 1b), suggesting the RELA part in the fusion protein somehow helped in stabilizing the binding. Furthermore, we selected genes that were bound by C11orf95^fus1^ and/or C11orf95-RELA^fus1^ at promoter regions, categorized them into three groups: C11orf95^fus1^ and C11orf95-RELA^fus1^ common binding genes, C11orf95^fus1^ unique binding genes and C11orf95-RELA^fus1^ unique binding genes. Gene ontology (GO) analysis showed C11orf95-RELA^fus1^ unique binding genes were specifically enriched in neuron projection morphogenesis, actin filament process and skeletal system morphogenesis, which are known to be associated with brain or brain disease development^3–6^. C11orf95^fus1^ and C11orf95-RELA^fus1^ common binding genes were enriched in cell cycle arrest, regulation of autophagy and regulation of mitochondrion organization. However, C11orf95^fus1^ unique binding genes were enriched in more fundamental biological processes such as translation, carbohydrate derivative biosynthetic process and asparagine N-linked glycosylation (Fig. 1c, Supplementary Table 1). In addition, we also mapped the transcription profiles in G16-3 and G16-4, as well as HEK293T control cells by RNA-seq to interrogate the transcriptional effects of C11orf95^fus1^ and C11orf95-RELA^fus1^. Sixty-six and 210 genes were identified to be up-regulated (p-value < 0.05, fold change >= 1.5) in G16-3 and G16-4 cells, respectively, comparing to the controls. In parallel, we compiled a list of 886 ST-EPN-RELA associated genes based on published data^1,7,8^. By comparing G16-3 and G16-4 up-regulated genes to ST-EPN-RELA associated genes, only three of G16-3 (C11orf95^fus1^) up-regulated genes overlapped with ST-EPN-RELA associated genes, while 67 of G16-4 (C11orf95-RELA^fus1^) up-regulated genes were in the ST-EPN-RELA associated gene list, including some well-known marker genes such as *L1CAM, CHD5* and *NOTCH1* (Extended Data Figs. 3a, b). All these evidences collectively indicate that C11orf95^fus1^ alone is not sufficient to drive transcriptomic dysregulation like C11orf95-RELA^fus1^.

In addition to HEK293T-based transgenic models, we also utilized another cell line, BXD-1425-EPN^7^ that was established from an orthotopic PDX originally derived from a ST-EPN-RELA tumor. Genotyping-sequencing and Western blot experiments with BXD-1425-EPN identified a novel fusion configuration containing the first three exons and a small fragment of the fourth exon of *C11orf95* gene, and *RELA* without the first two and half exons (Extended Data Fig 1c), here referred to as *C11orf95-RELA*^*1425*^. Despite the lack of fusion-specific antibody, the fact that unstimulated RELA is excluded from the nucleus^9^ while the fusion protein binds to DNA allowed us to use anti-RELA antibody to conduct ChIP-seq to pull down C11orf95-RELA^1425^ bound chromatin. As a result, RELA ChIP-seq in BXD-1425-EPN cells successfully identified 13,954 chromatin binding sites of C11orf95-RELA^1425^, with 5,338 commonly shared by C11orf95-RELA^fus1^ bindings in G16-4 cells. With the binding peak sequences of C11orf95^fus1^, C11orf95-RELA^fus1^ and C11orf-RELA^1425^, we implemented *de novo* motif discovery using MEME ChIP tools (http://meme-suite.org/tools/meme-chip). An identical GC-rich motif (GTGGCCCC) was readily recovered with top scores from all three sets of ChIP-seq peak sequences (Fig. 2a), supporting the validity of fusion protein pull-down by RELA antibody in BXD-1425-EPN cells and further confirming that the C11orf95^fus1^ fragment dictates DNA binding specificity of the fusion protein. As the motif consensus was repeatedly enriched by using different cells with different types of C11orf95-RELA fusion and different antibodies, the identified C11orf95 binding specificity is highly faithful. To establish GTGGCCCC as the *bona fide* DNA binding motif of C11orf95, we performed a reporter assay with 15 copies of GTGGCCCC and a minimum promoter cloned upstream of an EGFP reporter gene (Fig. 2b). As expected, C11orf95-RELA^fus1^ activated EGFP reporter expression harboring copies of the putative C11orf95 motif. In contrary, the unfused constituents RELA or C11orf95^fus1^ were unable to activate the reporter.

**Fig. 2.**
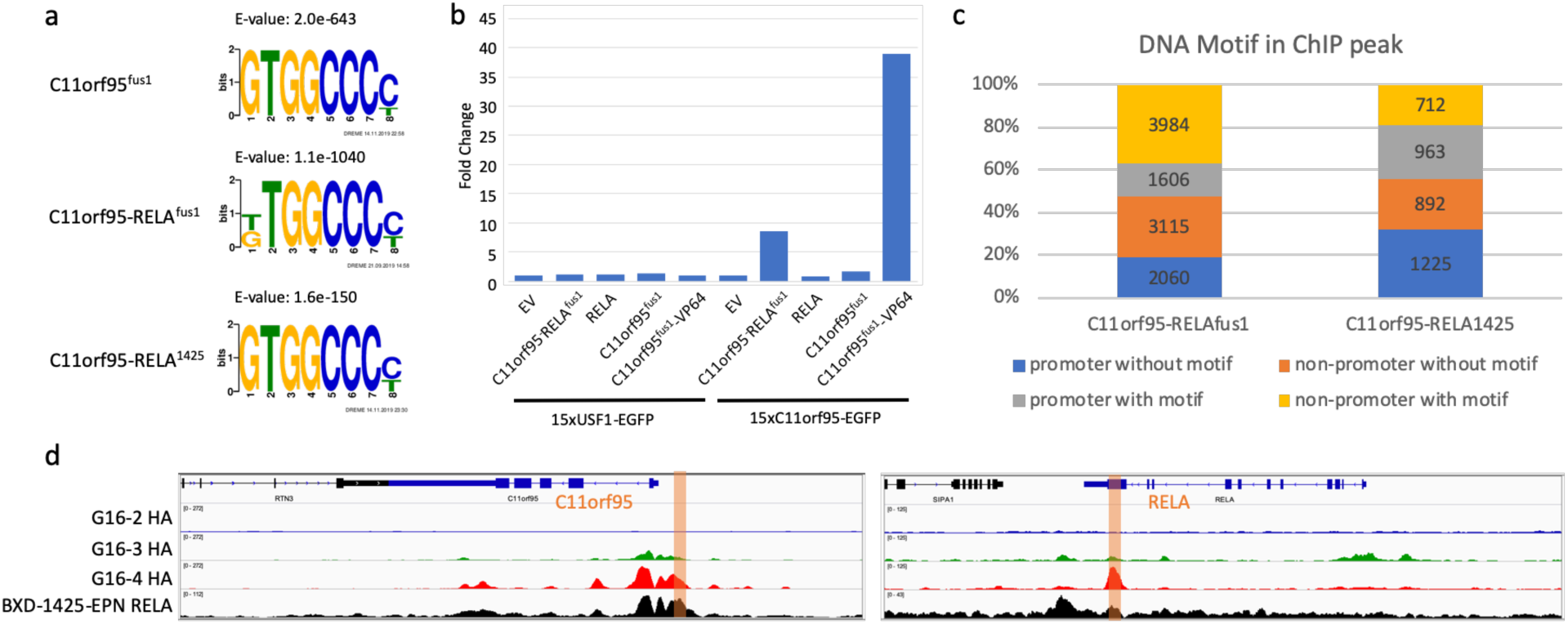
a. DNA binding motifs enriched for C11orf95^fus1^ and C11orf95-RELA^fus1^ and C11orf95-RELA^1425^ in G16-3, G16-4 and BXD-1425-EPN cells, respectively. b. Reporter assay. Reporter EGFP gene was linked to identified C11orf95 binding motif as well as USF1 motif as control. Empty vector (EV), RELA, C11orf95^fus1^ or activation domain VP64 fused C11orf95^fus1^ were expressed to test EGFP expression. EGFP signals were measured by flow cytometry. c. C11orf95 binding motif distribution in C11orf95-RELA peaks in G16-4 and BXD-1425-EPN cells. d. RELA, C11orf95^fus1^, C11orf95-RELA^fus1^ and C11orf95-RELA^1425^ binding profiles at *C11orf95* and *RELA* genes. The orange bars indicated the regions that had C11orf95 binding motifs.

Interestingly, supplementing C11orf95^fus1^ with a heterologous VP64 activation domain in the form of C11orf95^fus1^-VP64 fusion endowed transactivation activity on the reporter with 15 copies of the putative C11orf95 (GTGGCCCC) DNA binding motif but not on a control reporter harboring 15 copies of an unrelated USF1 motif. These results suggest that C11orf95^fus1^ binds to the GC-rich motif GTGGCCCC and may activate gene expression by hijacking the transcriptional activation domain of RELA, as in the case of the C11orf95-RELA fusion.

Among the top-score ChIP-seq peaks of C11orf95-RELA^fus1^ and C11orf95-RELA^1425^, approximately half of them have C11orf95 binding motif (Fig. 2c), implying a direct binding by the fusion. The no-motif peaks might be bound by a C11orf95-RELA-containing protein complex that used another pioneer subunit to initiate chromatin binding. Both motif and no-motif peaks were distributed at either promoter or non-promoter region with no significant bias (Fig. 2c). Intriguingly, ChIP-seq experiments in G16-4 and BXD-1425-EPN cells identified fusion binding peaks at C11orf95 promoter and RELA transcription end sites, with the C11orf95 DNA binding motif observed in both peaks (Fig. 2d), suggesting the possibility of auto-regulation.

## C11orf95-RELA modulates chromatin state and mediates chromatin interaction

We applied H3K27ac ChIP-seq to profile active chromatin regions in HEK293T transgenic cell models and BXD-1425-EPN to address the correlation between the fusion protein binding and chromatin state. At 3,792 fusion protein binding sites with top confidence in BXD-1425-EPN cells, strong H3K27ac signals were observed (Fig. 3a). Comparative analysis of genome-wide H3K27ac profiles between transgenic HEK293T cell models expressing fusion (G16-4) and RELA (G16-2) identified 441 differential H3K27ac peaks specific in G16-4 cells. These G16-4 specific H3K27ac peaks highly correlated with fusion protein bindings (Fig. 3b) with over 90% of them overlapping with fusion protein binding sites, implying the C11orf95-RELA was able to initiate chromatin state changes under certain circumstances. *CCND1* and *L1CAM*, two well-known marker genes of ST-EPN-RELA, exemplify two different chromatin state outcomes upon fusion protein binding. At *L1CAM* promoter region, fusion protein binding likely resulted in the deposition of H3K27ac marks (Figure 3c). In the contrary, *CCND1* promoter retained its active chromatin state regardless of the presence (in G16-4 and BXD-1425-EPN cells) or absence (G16-2 and G16-3 cells) of the fusion protein binding (Fig. 3c).

**Fig. 3.**
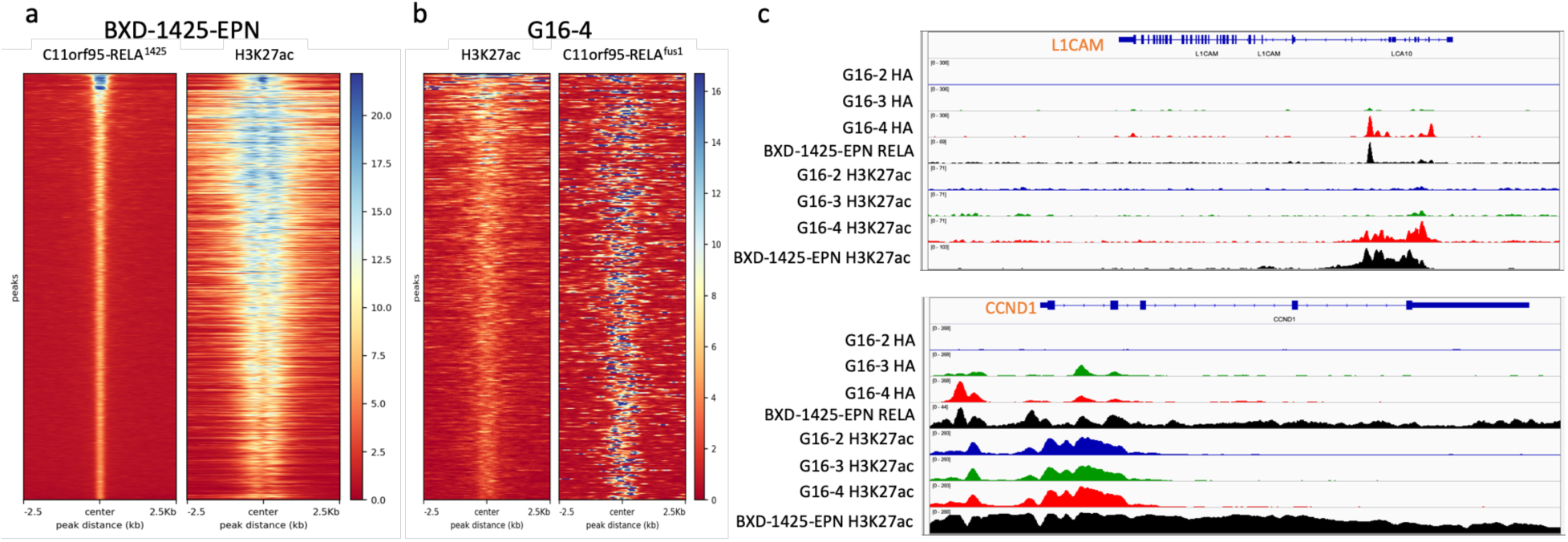
a. ChIP-seq read densities of C11orf95-RELA^1425^ and H3K27ac in BXD-1425-EPN cells. 3,792 top-score C11orf94-RELA^1425^ ChIP-seq peaks were used to plot the heatmap. The left side heatmap showed C11orf95-RELA^1425^ signals along a 5 kb region centering the peak summit. The right side heatmap showed H3K27ac signals along the same regions. b. Read densities of H3K27ac and C11orf95-RELA^fus1^ in G16-4 cells. 441 differential H3K27ac peaks in G16-4 compared to G16-2 cells were used to plot the heatmap. The left side heatmap showed H3K27ac signals along a 5 kb region centered with H3K27ac peak center. The right side heatmap showed C11orf95-RELA^fus1^ signals along the same regions. c. RELA, C11orf95^fus1^, C11orf95-RELA^fus1^ and C11orf95-RELA^1425^ as well as corresponding H3K27ac ChIP-seq profiles at ST-EPN-RELA marker genes *CCND1* and *L1CAM*.

As three-dimensional chromatin structure is known to be tightly associated with disease and cancer development, we next sought to identify changes in genome structure mediated by the fusion in ST-EPN-RELA. We utilized a sequencing-based chromatin capture approach HiC Chromatin Immunoprecipitation (HiChIP)^10^ to characterize genome-wide chromatin interactions and connect the protein binding peaks into loops and networks. We used anti-HA antibody to perform HiChIP of HA-tagged fusion protein in G16-4 cell lines. By further filtering with C11orf95-RELA^fus1^-HA ChIP-seq peaks, we identified a total of 32,669 high-confidence interactions. For BXD-1425-EPN cells, we performed H3K27ac HiChIP and identified 15,777 C11orf95-RELA1425 mediated interactions filtered by both H3K27ac and RELA ChIP-seq peaks. We mapped HiChIP identified chromatin interactions to the promoter regions of 886 ST-EPN-RELA associated genes mentioned above (Supplementary Table 2) to identify ST-EPN-RELA gene associated interactions. As a result, a total of 445 common interactions were identified in both G16-4 and BXD-1425-EPN cells. Interestingly, not all the 886 genes harbored chromatin interaction(s) to other locus/loci, but only 156 of them (Supplementary Table 3) reached out to one or more promoter or non-promoter (enhancer) regions. Since these ST-EPN-RELA gene associated chromatin interactions were directly mediated by the fusion protein, it is reasonable to speculate that the more interacting partners one gene had, the more regulatory events it was involved in, and the more important roles it played in ST-EPN-RELA. In other words, the gene that had more interactions to other genomic loci were more likely to be the hub of the fusion protein mediated network and more delicately regulated in fusion associated transcription, so that they might contribute more to ependymoma development. Hence, we counted the number of interactions each gene promoter is involved in and plotted a histogram ranking the 156 genes by their interaction counts (Fig. 4a). The top ranked genes included *CACNA1H, MAFG, NOTCH1, GPSM1, MXRA8, PYCR1, VWA1* and *RXRA*. Of note, *CACNA1H* is a calcium channel gene previously implicated and experimentally validated to be important in ST-EPN-RELA cell survival^11^, supporting the validity of our strategy in grading the significance of the genes. *NOTCH1* and *GPSM1* encode membrane proteins critical for NOTCH signaling and G-protein signaling, respectively, suggesting the involvement of these signaling pathways in ependymoma development in addition to NF-κB pathway that has received the most attention. To comprehensively nominate ependymoma-associated signaling pathways, we conducted a canonical pathway analysis with the genes that had more than one interacting chromatin site using Ingenuity Pathway Analysis (Qiagen) (Fig. 4b). Interestingly, the top enriched pathway was NOTCH signaling. Looking back at the ST-EPN-RELA gene list (Supplementary Table 2), 15 Notch signaling genes stood out, *ADAM12, CCND1, DLK1, DVL1, DVL2, HDAC1, HES1, HES5, JAG1, JAG2, LFNG, LNX1, MFAP2, NOTCH1* and *PSENEN*, further demonstrating Notch signaling might play a significant role in ST-EPN-RELA oncogenesis^12^.

**Fig. 4.**
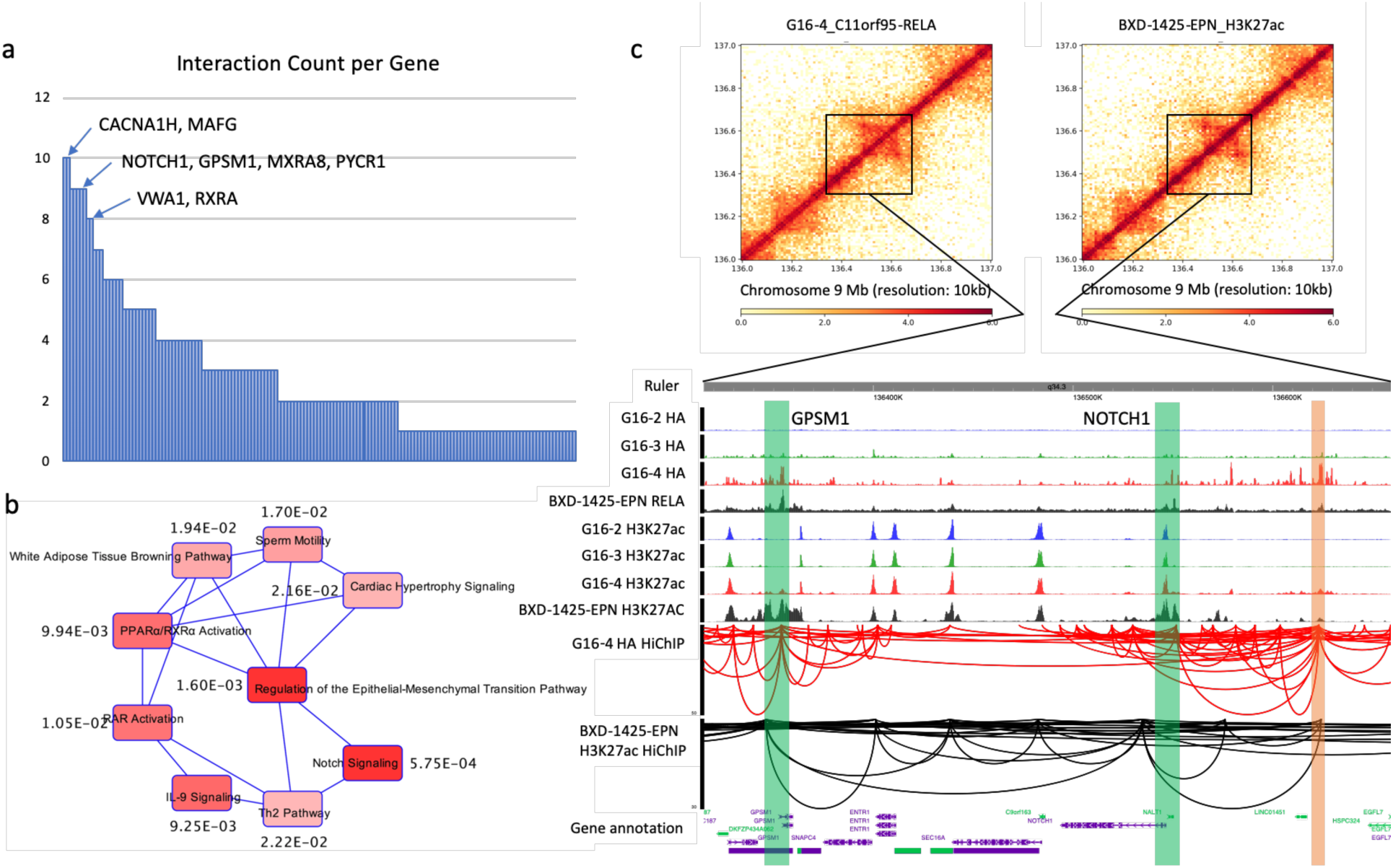
a. Histogram of interaction counts per gene. The interactions coming out from the 156 identified gene promoters were counted. b. Canonical pathway analysis of 102 genes that had more than one interaction. The numbers by the pathways are p-values. c. Chromatin interactions captured by HiChIP along *NOTCH1* and *GPSM1* genes. The heatmaps showed a 1 Mb region of interactions along *NOTCH1* and *GPSM1* genes. The browser view showed the zoom-in details of the black box indicating region in the heatmaps, including ChIP-seq profiles and HiChIP interactions. The green bars indicate *NOTCH1* and *GPSM1* promoters. The orange bar indicates the enhancer.

As we mentioned above, the fusion protein not only mediated promoter-promoter interactions but also enhancer-promoter interactions. To this end, we again made use of the 445 ST-EPN-RELA gene associated chromatin interactions identified in both G16-4 and BXD-1425-EPN cells, but focusing on the 115 non-promoter (putative enhancer) regions (Supplementary Table 4). We then counted the number of gene promoters connecting to these 115 enhancers. Although the majority of the enhancers had only one interacting promoter, some of them interacted with two to seven promoters (Extended data Fig. 4a). The enhancer with the highest number of promoter interactions is located in the intron of *ASPSCR1* gene at chromosome 17, connecting with *ASPSCR1, DCXR, MAFG, MAFG-AS1, NPB, PYCR1* and *RAC3* genes (Extended data Fig. 4b). More impressively, another distal enhancer had robust interactions with promoters of *NOTCH1* and *GPSM1* genes in both G16-4 and BXD-1425-EPN cells. Fusion protein binding was intensive and resulted in H3K27ac deposition at the enhancer region in G16-4 and BXD-1425-EPN cells (Figure 4c). This suggests a potential regulatory scenario in which the fusion protein induces epigenetic changes at an enhancer and mediates chromatin reorganization with key genes of two distinct arms of signaling, specifically the Notch and G-protein signaling pathways, to initiate genetic programs underlying ST-EPN-RELA pathogenesis.

Informed by our study of epigenetic reprogramming driven by C11orf95-RELA fusion in HEK293T cell models and ependymoma cell line, we gained important knowledge on the molecular mechanisms underlying tumorigenesis driven by a single mutation of C11orf95-RELA fusion^1,13^. Although the overexpression of C11orf95^fus1^ or RELA alone in HEK293T models were not sufficient to cause ependymoma-like transcriptomic transformation, C11orf95 portion in the fusion plays a more critical role than merely serving as a shuttle for the entry of RELA to the nucleus. Our data shows that C11orf95 recognizing a specific DNA motif and regulating gene expression by hijacking the activation domain of RELA. Since the first discovery of C11orf95-RELA mutation in ST-EPN^1^ (Parker 2014), NF-κB pathway caught the most attention in tumorigenesis signaling. However, it is not the only pathway involved in ST-EPN-RELA (Ozawa 2018). In addition, no significant change of C11orf95-RELA ependymoma cell viability was observed when they were treated with NF-κB inhibitors^14^, further implying that NF-κB might not be the major player in driving the malignant phenotype of ST-EPN-RELA. Our study showed that Notch signaling ranked among the top pathways suggested by ST-EPN-RELA genome and chromatin structure analyses as a number of Notch genes were identified as chromatin hubs harboring more than one distal interacting partners. Therefore, Notch inhibitors or a combination of Notch and NF-κB inhibitors could be better candidates for ST-EPN-RELA treatment. Other than the direct binding to gene promoters, C11orf95-RELA also binds to regulatory elements such as enhancers. It is interesting to know that enhancer could be multi-functioning in regulating multiple ST-EPN-RELA genes, which implies that a potential gene therapy strategy is to target the enhancers.

## Methods

### Cell culture

HEK293T and its derived G16-2, G16-3, G16-4 cells as well as BXD-1425-EPN cells were cultured in Dulbecco’s modified Eagle’s medium (Sigma) with 10% fetal bovine serum (Lonza), 4% Glutamax (Gibco), 1% Sodium Pyruvate (Gibco) and penicillin-streptomycin (Gibco). Incubator conditions were 37 °C and 5% CO2. Protein overexpression in G16-2, G16-3 and G16-4 cells were induced by 1 μg/ml doxycycline treatment for two days.

### Virus production

A viral packaging mix of pLP1, pLP2, and VSV-G were co-transfected with each lentiviral vector (pCW-RELA-HA, pCW-C11orf95^fus1^-HA, or pCW-C11orf95-RELA^fus1^-HA) into Lenti-X 293T cells (Clontech), seeded the day before in 6-well plates at a concentration of 1.2×10^6^ cells per well, using Lipofectamine 3000 (ThermoFisher). Media was changed 6 hours after transfection then incubated overnight. Twenty-eight hours post transfection, the media supernatant containing virus was filtered using 45 μm PES filters then stored at −80 °C until use.

### Cell Line generation

To make G16-2, G16-3, and G16-4 cell lines, the day prior to transduction, HEK293T cells were seeded into 12-well plates at a density of 1.5×10^5^ cells per well. Prior to transduction, media was changed to that containing 10 μg/mL polybrene, 1 mL per well. 250 μL of each respective virus was added to each well and incubated overnight. Media was changed 24-hour post-transduction. Four days post-transduction, cells were split. Five days post-transduction, media with antibiotics (2 μg/mL Puromycin) was added to each respective well of one replicate plate. Antibiotic selection continued for at least 2 weeks before use in downstream experiments.

### Cloning

ORF encoding C11orf95^fus1^ was obtained as IDT gBlock while HEK293T cDNA served as template for amplifying RELA sequence. PCR and SLIC cloning were used to insert ORF encoding C11orf95^fus1^, C11orf95-RELA^fus1^, RELA with C-terminal HA tag into Gateway donor vector (pCR8/GW/TOPO, Invitrogen). Then LR clonase II reactions were used to shuttle ORFs into pCW-DEST (lentiviral Dox-inducible expression) derived from pCW-Cas9 (Addgene # 50661), and pmax-DEST (transient constitutive expression, Addgene # 48222) vectors, generating pCW-C11orf95^fus1^-HA, or pCW-C11orf95-RELA^fus1^-HA, pCW-RELA-HA lentiviral Dox-inducible plasmids and pmax-C11orf95^fus1^-HA, pmax-C11orf95-RELA^fus1^-HA, pmax-RELA-HA transient expression plasmids. pmax-Clover and pmax-C11orf95-Clover, and pmax-C11orf95^fus1^-VP64, pmax-mScarlet-H2A, plasmids were generated with a combination of PCR, restriction-ligation, SLIC and LR Clonase II reactions. GFP-reporter plasmids, 15xUSF1-minCMV-EGFP and 15xC11orf95-minCMV-EGFP, were generated by ligation of annealed oligos containing USF1 or C11orf95 binding sites into minCMV-EGFP plasmid with non-palindromic overhang sites via digestion with BsaI site, subsequently screened for the number of binding sites by Sanger sequencing.

### Transfection and imaging

Cells were seeded into 12-well plates at a density of 1.5×10^5^ cells per well the day before transfection. For imaging experiments, 300 ng of pmax-C11orf95^fus1^-Clover or pmax-Clover (unfused control) was co-transfected with 100 ng of pmax-mScarlet-H2A (nuclear marker) using 1.5 µL Attractene transfection reagent (Qiagen). Microscopy images were taken with the iRiS Digital Cell Imaging System (Logos Biosystems).

### Flow cytometry

GFP-reporter plasmid DNA (300ng) containing 15xUSF1 binding sites (15xUSF1-minCMV-EGFP) or 15xC11orf95 binding sites (15xC11orf95-minCMV-EGFP) was co-transfected with 300 ng of plasmid expressing test transcription factor constructs (EmptyVector, pmax-C11orf95^fus1^, pmax-C11orf95-RELA^fus1^, pmax-RELA, pmax-C11orf95^fus1^-VP64) using 1.5 µL Lipofectamine 3000 (ThermoFisher). 48 hours after transfection, cells were trypsinized, suspended in media then analyzed on a LSRFortessa X-20 flow cytometer (BD Bioscience). Fifty thousand events were collected each run.

### Western blot

Western blot was performed using general protocol from Bio-Rad. Primary antibodies used were HA antibody (Cell Signaling Technology 3724S), RELA antibody (Abcam ab32536), and β-actin antibody (Cell Signaling Technology 4970S).

### Genotyping

DNA was extracted using DNeasy Blood & Tissue Kit (Qiagen), followed by genotyping PCR using Phusion PCR protocol (NEB) with primer pair of forward: CCTGCACCTGGACGACAT and reverse: TTGGTGGTATCTGTGCTCCTC. PCR amplicons were sent for Sanger sequencing, and analyzed by ApE.

### Gene ontology analysis

Gene ontology analysis was done with Metascape (http://metascape.org/).

### DNA motif analysis

ChIP-seq peak sequences were uploaded and analyzed by MEME-ChIP (http://meme-suite.org/tools/meme-chip). Motifs were found *de novo* by DREAM. Sequences that had specific motif were reported by FIMO.

### RNA-seq and analysis

RNA was extracted using RNeasy Mini Kit (Qiagen), then sent to Genewiz for library construction and sequencing by Illumina Nextseq sequencer. Each sample had two biological replicates. RNA-seq data was quantified using Salmon (version 0.8.0), followed by differential expression analysis using DEseq2. The cutoff for identifying differentially expressed genes were set as q-value <0.05 and fold change >=1.5.

### ChIP-seq and analysis

ChIP was performed using iDeal ChIP-seq kit for Transcription Factors (Diagenode) and protocol provided. HA (Cell Signaling Technology 3724S), RELA (Abcam ab32536) and H3K27ac (Cell Signaling Technology 8173S) antibodies were used for chromatin pull-down. Libraries were constructed using KAPA Hyper Prep Kits (Roche) and NEBNext^®^ Multiplex Oligos for Illumina^®^ (NEB) for Illumina Nextseq sequencing. Each sample had two biological replicates. The reproducibility between replicates was evaluated by Pearson correlation coefficient of read counts using deepTools (version 3.3.0) which was all greater than 0.95. Replicates were then combined for peak calling. Sequencing data was mapped to hg38 reference genome by Bowtie2 (version 2.3.1). Sequences with mapping quality score greater than ten were accepted for the following analysis. Peaks were called by MACS2 (version 2.1.0.20151222) using “-q 0.01 --nomodel --extsize 200” parameter for transcription factor ChIP-seq, and using “--broad -q 0.05, --nomodel --extsize 220” for histone mark ChIP-seq. For peaks that had at least 1 bp overlap, we considered them as overlapped peaks. The peak was considered to locate at gene promoter when it had at least 1 bp overlap with the promoter region. Promoter region was defined as a 5 kb span extending 2.5 kb at both sides from the transcription start site (TSS). Differential H3K27ac peaks were called by MACS2 bdgdiff subcommand. Read density heatmap of ChIP-seq peaks were computed and plotted using deepTools (version 3.3.0). Data was viewed by Integrative Genomics Viewer (IGV).

### HiChIP and analysis

HiChIP was performed following protocol published by Mumbach et al. 2017, with minor modifications. HA (Cell Signaling Technology 3724S) and H3K27ac (Cell Signaling Technology 8173S) antibodies were used. Libraries were constructed using KAPA Hyper Prep Kits (Roche) and NEBNext^®^ Multiplex Oligos for Illumina^®^ (NEB) for Illumina Novaseq sequencing. Each sample had two biological replicates. Replicate data were combined for analysis. Raw data was processed by HiC-Pro^15^ using hg38 reference genome. Chromatin interactions were called using hichipper^16^ by providing certain ChIP-seq peaks and HiC-Pro outputs. Interaction heatmap was plotted by HiCPlotter^17^ at 10 kb resolution. Interactions associated with specific genes were filtered by interaction count greater than one, then visualized on WashU Epigenome Browser, or processed for enhancer-promoter analysis.

### Canonical pathway analysis

Canonical pathway analysis was done by Ingenuity Pathway Analysis (IPA) using “Core Analysis” function.

## Acknowledgements

We would like to thank members of the C.C.L. laboratory at the Jackson Laboratory for Genomic Medicine for helpful discussions and Amanda Lazarus for administrative support. This work was partially supported by funding from the Chenevert Foundation (C.C.L.) and the Martin J. Gavin Endowment at Connecticut Children’s Medical Center (C.C.L.), National Cancer Institute P30CA034196 (to A.W.C.) and the National Human Genome Research Institute R01-HG009900 (to A.W.C.).

## Author contributions

C.C.L. and A.W.C. conceived the project and led the studies; J.J.Z. A.W.C and C.C.L designed the experiments. J.J.Z., N.J. and A.W.C. carried out the experiments; X-N.L. developed the orthotopic PDX model BXD-1425-EPN and established the corresponding cell line; J.J.Z., A.W.C. and C.C.L. analyzed and interpreted the results, J.J.Z., A.W.C and C.C.L. wrote the manuscript and all authors reviewed the manuscript.

## Competing interest declaration

The authors declare that they have no competing interests.

## Extended data

**Extended Data Fig. 1.**
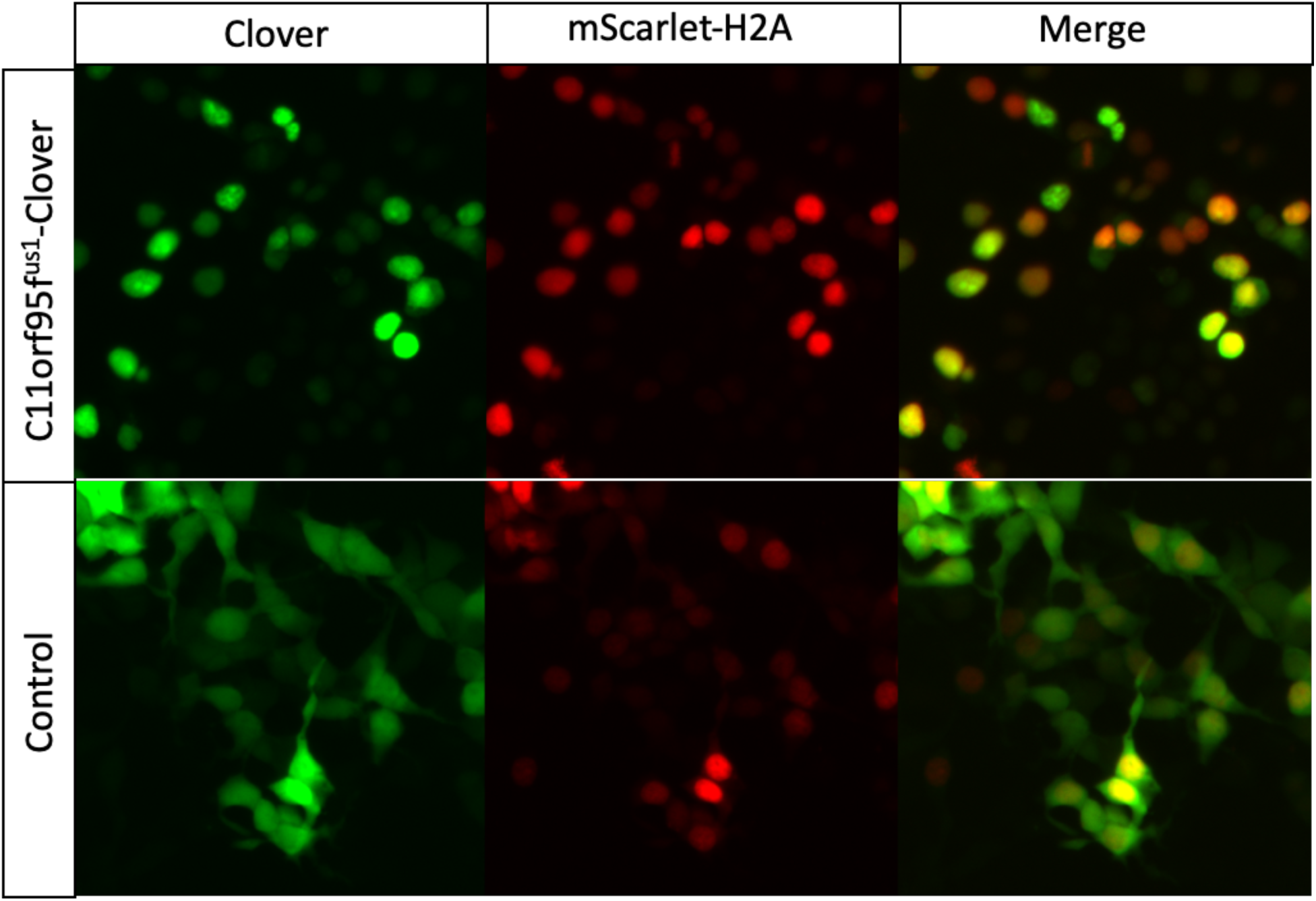
Colocalization of C11orf95^fus1^ with cell nucleus. Clover indicate C11orf95^fus1^ fused or unfused clover protein in C11orf95^fus1^-clover or control cells, respectively. mScarlet-H2A indicate cell nucleus.

**Extended Data Fig. 2.**
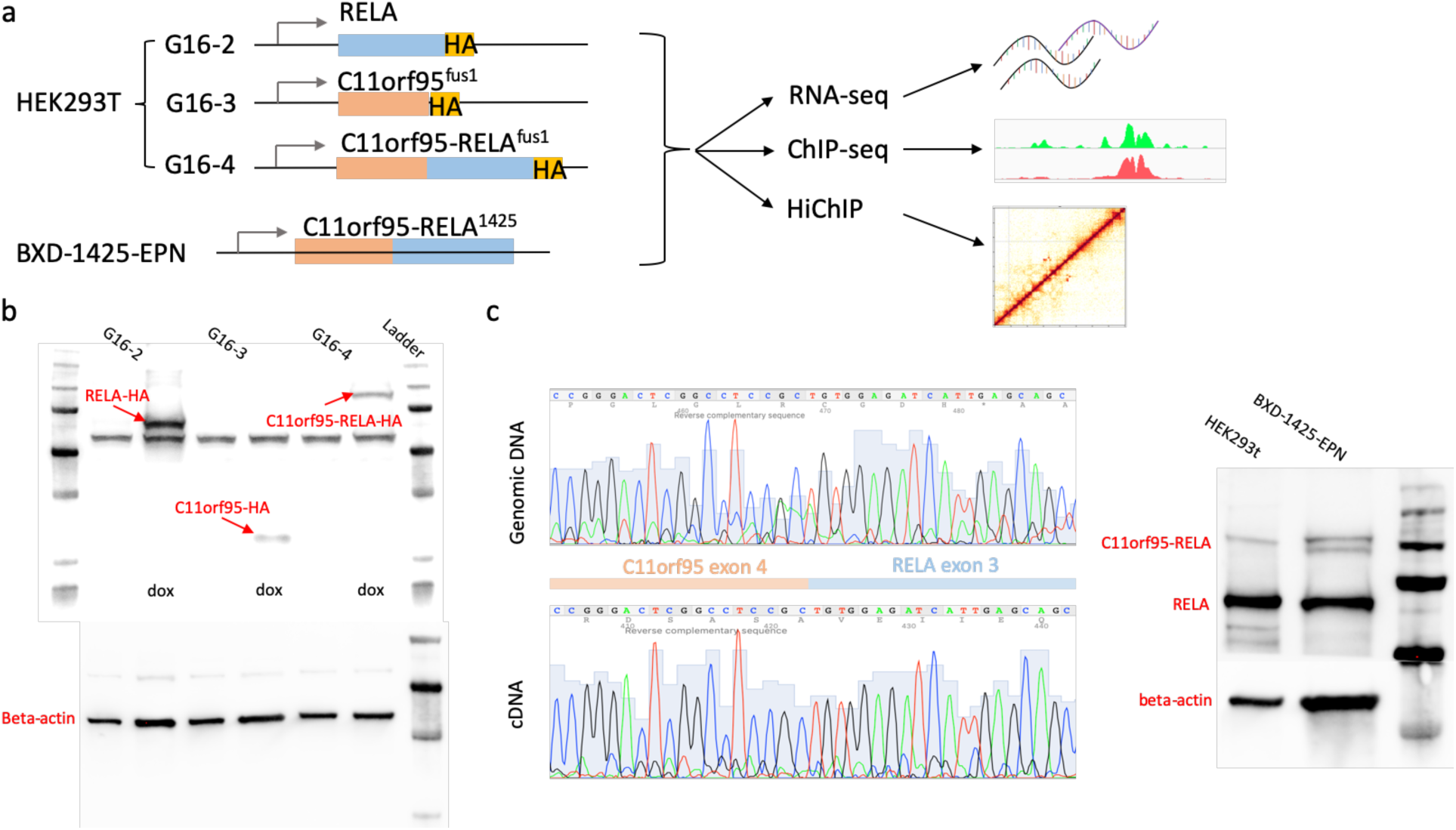
a. Flow chart of different cell lines used for different sequencing mappings. b. Western blot for RELA, C11orf95fus1 and C11orf95-RELAfus1 overexpression in G16-2, G16-3, G16-4 cells. Blue arrow points to the overexpressed protein marked out. Each cell line has two lanes. One lane is for doxycycline-treated cells, the other is for non-treated cells as control. Beta-actin is used as internal protein control for each cell type. c. Confirmation of fusion type in BXD-1425-EPN cells with genotyping and western blot.

**Extended Data Fig. 3.**
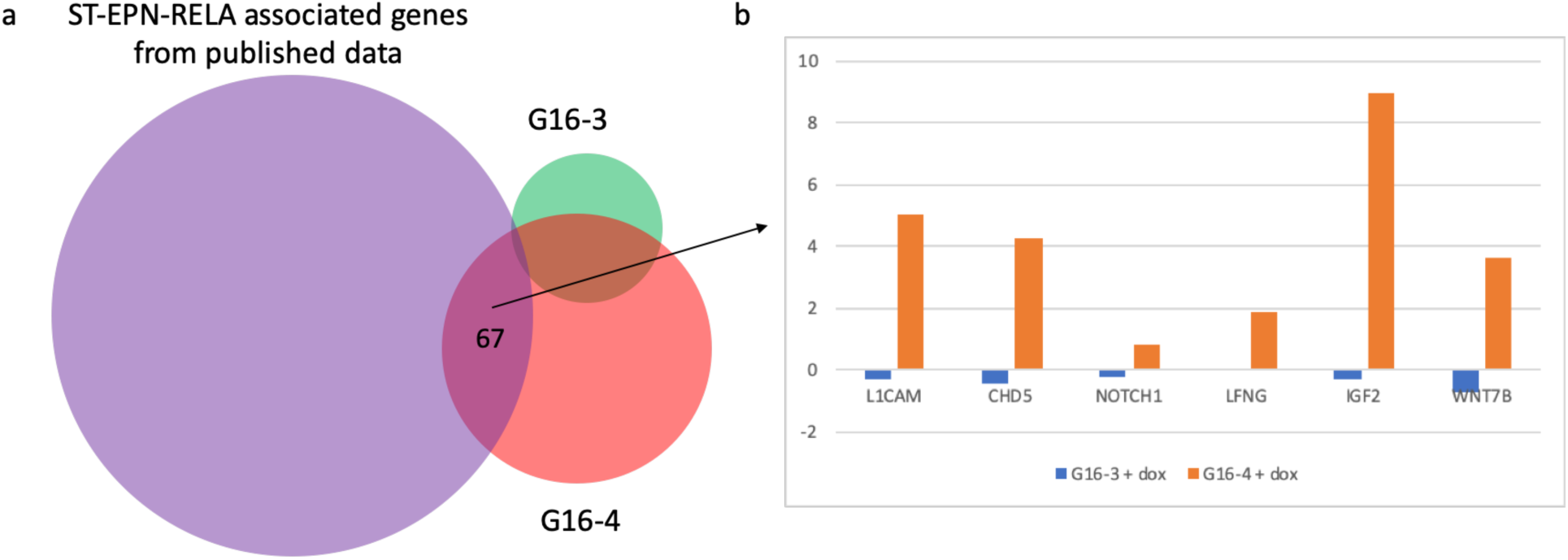
a. Venn diagram of ST-EPN-RELA associated genes from published data (supplementary Table 2) (purple) and upregulated genes in G16-3 (green) and G16-4 cells (red). Upregulated genes in G16-3 and G16-4 cells were identified by RNA-seq by comparing to HEK293T cells. There were very few overlaps of G16-3 upregulated genes and published ST-EPN-RELA genes, but 67 including well-known ST-EPN-RELA associated genes (*L1CAM, CHD5, NOTCH1, LFNG, IGF2* and *WNT7B*) were upregulated in G16-4 cells. The Venn diagram was not drawn to scale. b. Expression of well-known ST-EPN-RELA associated genes in G16-3 and G16-4 cells.

**Extended Data Fig. 4.**
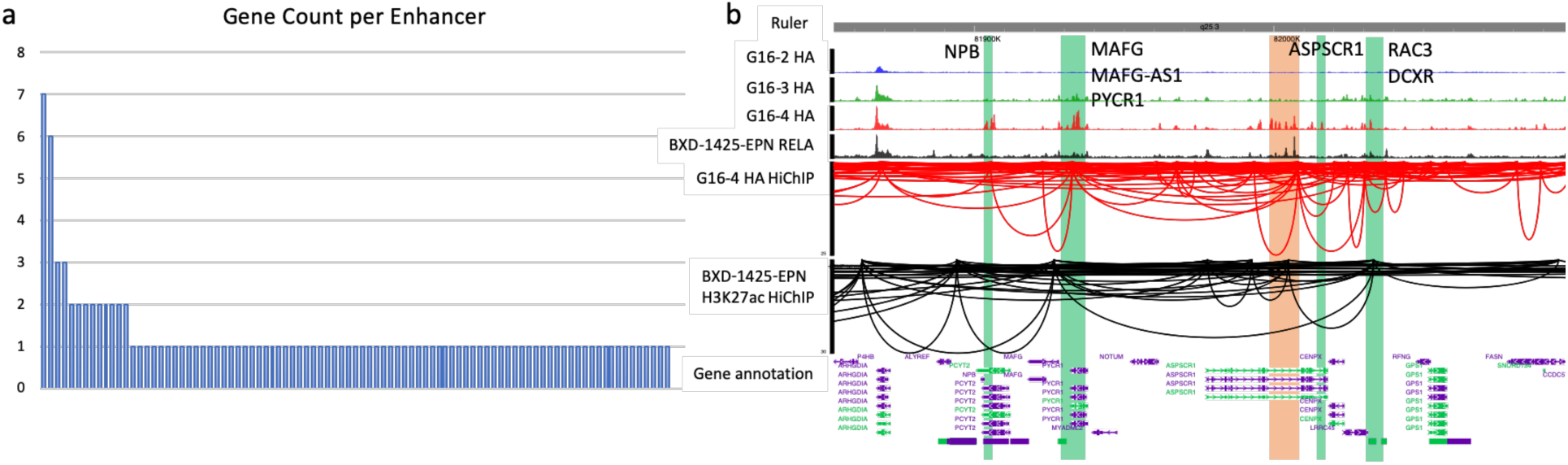
a. Histogram of interaction counts per enhancer. 115 ST-EPN-RELA associated enhancer-promoter interactions were analyzed. The counts indicate the number of ST-EPN-RELA genes that are regulated by the same enhancer. b. Browser view of the enhancer that targets seven ST-EPN-RELA genes. The green bars indicate gene promoters. The orange bar indicates the enhancer.

## Notes

### Competing Interest Statement

The authors have declared no competing interest.

